# Structure-based Design of Chimeric Influenza Hemagglutinins to Elicit Cross-group Immunity

**DOI:** 10.1101/2024.12.17.628867

**Authors:** Karla M. Castro, Reyhaneh Ayardulabi, Sarah Wehrle, Hongrui Cui, Sandrine Georgeon, Joseph Schmidt, Shuhao Xiao, Nishat Seraj, Wayne Harshbarger, Corey P. Mallett, Ventzislav Vassilev, Xavier Saelens, Bruno E. Correia

## Abstract

Antigenic variability among influenza virus strains poses a significant challenge to developing broadly protective, long-lasting vaccines. Current annual vaccines target specific strains, requiring accurate prediction for effective neutralization. Despite sequence diversity across phylogenetic groups, the hemagglutinin (HA) head domain’s structure remains highly conserved. Utilizing this conservation, we designed cross-group chimeric HAs that combine antigenic surfaces from distant strains. By structure-guided transplantation of receptor-binding site (RBS) residues, we displayed an H3 RBS on an H1 HA scaffold. These chimeric immunogens elicit cross-group polyclonal responses capable of neutralizing both base and distal strains. Additionally, the chimeras integrate heterotrimeric immunogens, enhancing modular vaccine design. This approach enables the inclusion of diverse strain segments to generate broad polyclonal responses. In the future, such modular immunogens may serve as tools for evaluating immunodominance and refining immunization strategies, offering potential to bridge and enhance immune responses in individuals with pre-existing immunity. This strategy holds promise for advancing universal influenza vaccine development.

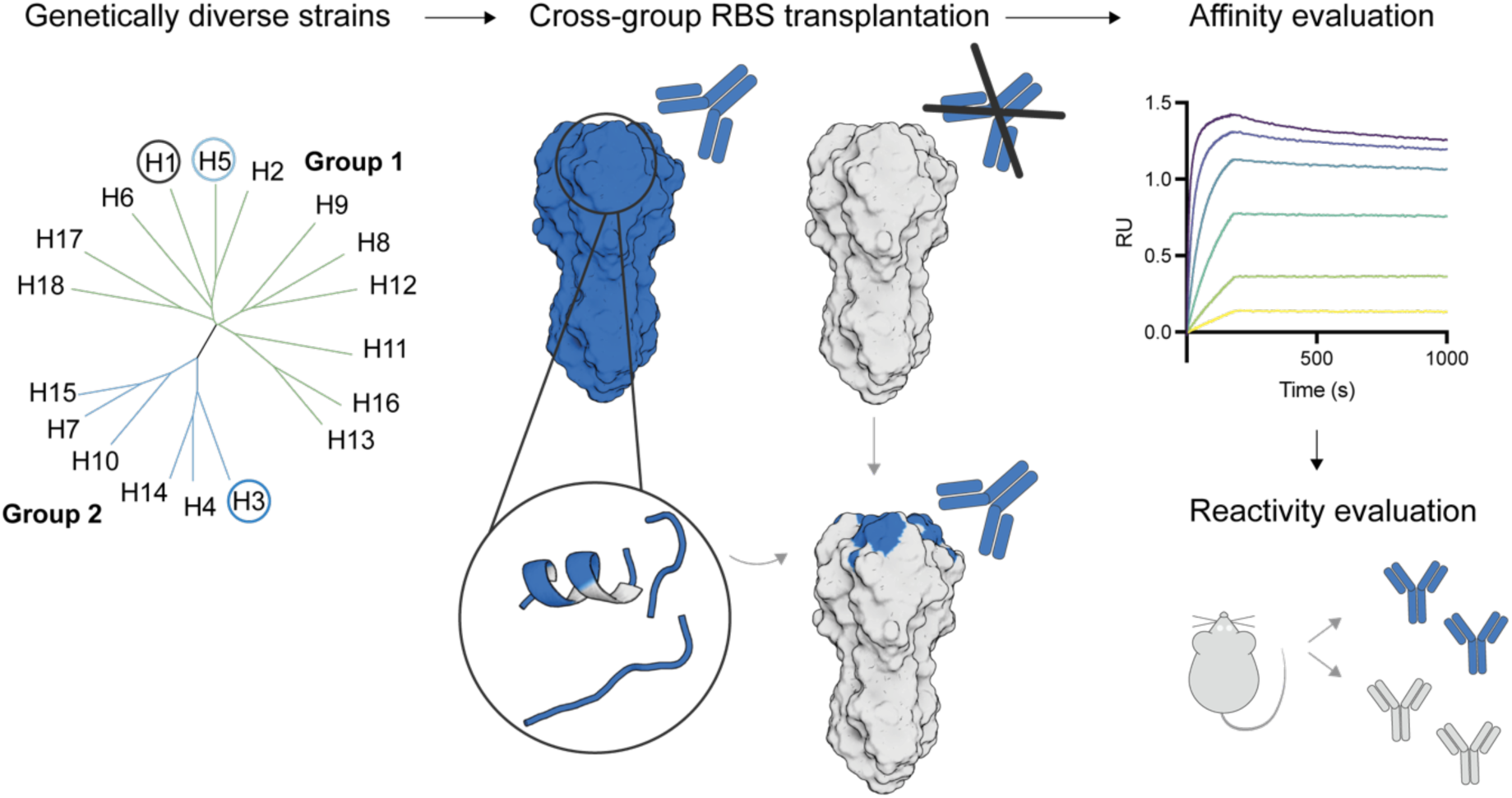
Graphical abstract: Overview of cross-group RBS transplantation approach. Phylogenetically diverse HA strains can be incorporated into chimeric immunogens by RBS transplantation. The chimera are evaluated for cross-reactivity to subtype-specific antibodies and the ability to elicit neutralizing antibodies to multiple strains.

## Introduction

Influenza is a major public health burden with vaccine formulation requiring yearly intervention based on annual predictions of circulating strains^1^. The current seasonal influenza vaccine includes hemagglutinin derived from H1N1, H3N2 and one or two lineages of influenza B. In addition, there are numerous animal influenza A virus subtypes, some of which are zoonotic such as highly pathogenic avian H5N1 viruses^2–4^. Current vaccines are based on the major viral glycoprotein hemagglutinin (HA), which consists of a receptor binding site (RBS) as a major antigenic site^5–8^.The head domain of HA, however, is antigenically variable and immunodominant. Therefore, to try to elicit broader immune responses against HA, several approaches have been proposed. One approach aimed to elicit broad H1 reactivity by presenting several H1 head domains on a nanoparticle, to boost subdominant cross-reactive B-cell responses and disfavour strain-specific responses^9^. An alternative approach was based on a consensus sequence approach to induce a broader range of antibodies against a particular subtype^10,11^. Other methods including beads-on-a-string (BOAS) and multivalent virus-like particle presentation aimed to present multiple strains as individual trimers together on one particle to induce broad protection ^12–14^. Recently, genetically fused HAs of three strains from group 1 were successfully presented on one trimer^15^. However, broadening the overall immune response by incorporating cross-group sequences on single trimer immunogens has not yet been demonstrated, likely due to structural incompatibilities of group 1 and group 2 sequences.

One approach to do so involves RBS sequence exchange. HA sequence replacements have successfully incorporated inter-subtype exchanges aimed to boost and broaden RBS-targeted antibody elicitation^16,17^. Furthermore, we hypothesised that the structural similarity in the HA head domain would be sufficient to tolerate changes required for sequence replacement of pandemic-relevant subtypes and that a stable trimeric immunogen with mixed head domains from different subtype HAs would be able to elicit a broader polyclonal response to cross-group HA subtypes towards expanding vaccine breadth in contrast to a vaccine cocktail. Here we design a cross-subtype HA chimera by structure-informed sequence exchange aimed to elicit an immune response to multiple HA subtypes of interest within and across groups which can also be readily incorporated as chimeric heterotrimers. In mice, the chimeric immunogens can elicit polyclonal cross-group immune responses. This study supports the use of cross-group chimeras as an efficient approach for multi-strain presentation in an influenza vaccine and an avenue for modularly incorporating vastly diverse HA subtypes on one trimer immunogen to increase the protective breadth of synthetic influenza immunogens.

## Results

### Cross-group chimeric RBS by sequence replacement

The natural receptor that engages HA, sialic acid, binds through highly conserved residues in the core of the RBS, while most known neutralising antibodies bind through less conserved residues in the surrounding 4 segments of the RBS ring: the 190-helix, 220-loop, 150-loop, and the 150-loop (Fig. 1A)^7,18,19^. We first explored the structural similarity of the HA head domain with closely related subtypes as well as distant cross-group subtypes. The four major RBS segments (Fig. 1A) show low sequence conservation between subtypes (Fig. 1B). However, the HA head domain shows remarkable structural similarity even across groups, particularly at the RBS (Fig. 1C). We specifically compared H1 of A/Solomon Islands/3/2006 (SI06) to H5 of A/Vietnam/1194/2004 and H3 of A/Hong Kong/1/1968, referred to in this paper as H1SI06, H5VN04, and H3HK68, respectively. HAs from these viruses have been structurally characterised in complex with RBS-specific antibodies. While H1SI06 was additionally selected for robust structural stability, H5VN04 and H3HK68 have been extensively structurally characterised in the presence of a number of neutralising antibodies^7,18,19^.

**Figure 1:**
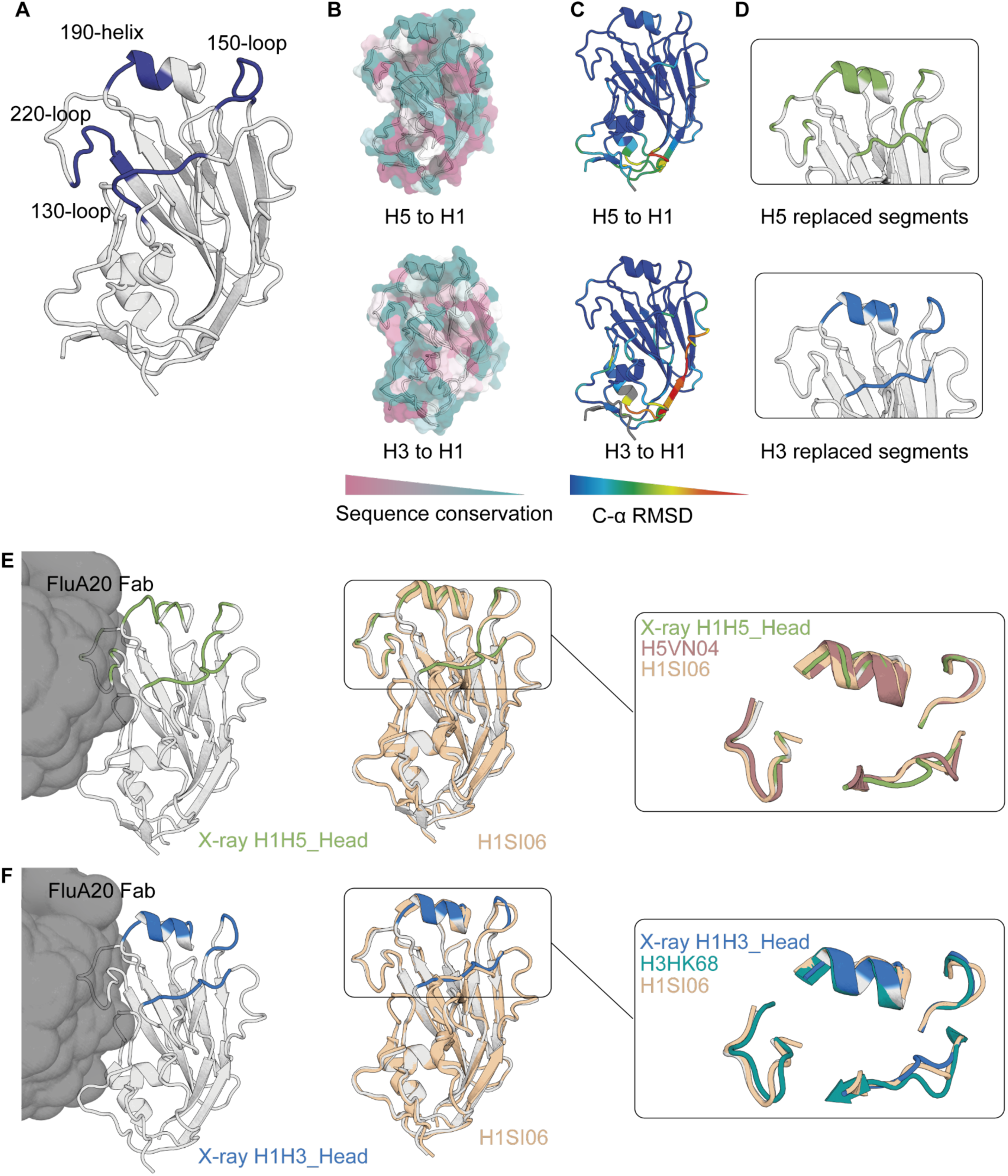
RBS is structurally conserved between strains. A) Structure of the H1SI06 hemagglutinin (PDB: 6OC3) epitope highlighting the four major RBS regions. B) Sequence conservation visualisation between H1SI06 and H5VN04, and H1SI06 and H3HK68. C) Structural conservation between H1SI06 and H5VN04, and H1SI06 and H3HK68, illustrated as RMSD_CA_ distance. D) AF2 predicted model of H1H5_head and H1H3_Head. Replaced segments from H5VN04 and H3HK68 are highlighted in green and blue, respectively. E-F) Crystal structure of the H1H5_Head and H1H3_Head chimera in complex with FluA20 Fab, where the sequence transplanted segments are coloured green and blue, respectively. The right panel shows the chimera (grey) overlaid on the H1SI06 WT (wheat) and zoomed in on the main RBS segments showing structural similarity of 3 segments and variability of loop 130 compared to H5VN04 WT apo and H3HK68 WT in complex with FluA20 (PDB: 4BGW and 6OCB, respectively).

We hypothesised that the structural similarity could be leveraged to replace the H1 RBS with other subtypes to present defined antigenic sites from more than one strain. We first exchanged cross-subtype residues within group 1. We replaced RBS residues that are surface exposed and critical for interaction with a subtype-specific antibody of H1SI06 with H5VN05^18^. The resulting H1H5_Head chimera presented the RBS of H5VN04 in a background of H1SI06, the two strains originally showing 40.6% sequence identity of the four RBS segments (Fig. 1D). We used binding of the HA trimer interface-specific antibody FluA20^20^ as an indicator of the adoption of native-like conformation, as this antibody relies on main chain interactions in the 220-loop provided the RBS is in native conformation. HA Head constructs H1H5_Head showed 62.6 nM apparent affinity for FluA20 IgG compared to 12 nM observed with H1SI06_Head wildtype while H1H5_Head showed low apparent affinity for a H5 RBS specific antibody, the kinetics could not be fit (Supplementary Fig. 1A-C). We then sought to assess the increased complexity of replacing cross-group residues from H3HK68 (H1H3_Head) where the sequence similarity of the RBS segments falls to 26.7% (Fig. 1D). While we observed 13.7 nM apparent affinity for FluA20 IgG, we could not detect binding of H1H3_Head to an H3 RBS-specific antibody (Supplementary Fig. 1D) suggesting the chimera adopted an overall Head conformation similar to native H1SI06, the RBS of H1H3_Head may not present full mimicry to H3 RBS in solution.

We sought to further investigate the structural integrity of the head domain as a result of cross-group residues replacement. While H1H5_Head showed a similar CD profile to the H1SI06 WT, we observed a decrease in Tm from 61°C in the H1SI06 to 40°C in the H1H5_Head (Supplementary Fig. 2A-B). The H1H3_Head showed a mixed coiled coil profile suggesting non-globularity in solution (Supplementary Fig. 2C). Yet, upon solving the crystal structure of both H1H5_Head and H1H3_Head in complex with FluA20 IgG, the transplanted RBS segments largely retain the structure of H1SI06 with the exception of the 130-loop not representative of either the H1SI06 WT base or the RBS replaced strain (Fig. 1E-1F). While this loop does not play a role in binding H5-specific antibody 13D4 IgG^18^, there are at least three major contacts in the 130-loop for binding to H3-specific RBS targeting antibody F045-92 IgG^7^ which may be flexible in FluA20 binding or alone in solution compared to when bound to the RBS-targeting antibody (Supplementary Fig. 3). Taken together, these data show that replacement of the RBS segments with cross-subtype and distant cross-group residues largely conserves RBS structure upon the stabilisation by FluA20 binding and may benefit from internal structural stabilisation to accurately present the RBS in solution.

### Trimeric constructs improve stability and antigenic profile

Following the observation that the HA head chimera can be stabilised when complexed to FluA20, we reasoned that full length homotrimer constructs would similarly enhance the stability of the head domain and consequently contribute to accurate presentation of the engineered RBS. Previous studies have shown modularity of HA head and stem domains^21,22^. We similarly incorporated the H1H5_Head and H1H3_Head with a H1SI06 stem sequence using a previously described approach to combine heterotypic head-stem constructs^23^ (H1H5_FL and H1H3_FL, and H1H5_FL2 and H1H3_FL2) (Fig. 2A). The designs H1H5_FL and H1H3_FL bound with an apparent K_D_ >1 μM and 441 nM, to H5-RBS-specific 13D4 IgG^18^ and H3-RBS-specific F045-92 IgG^7^, respectively (Fig. 2C-D). The affinities of H5VN04 and H3HK68 were 41 nM and 10 nM, respectively. We further improved the transplanted segments to better recapitulate subtype-specific antibody RBS binding by including additional epitope residues as observed in the H5-13D4 (PDB:6A0Z) and H3-F045-92 (PDB: 4O58) complexed crystal structures; residues were reverted to H1SI06 sequences for positions not directly contributing to subtype-specific antibody binding. Additionally, where the H5 or H3 subtype consensus residue identity differed from the specific strain residues of H5VN04 or H3HK68, consensus residue identities were incorporated (Fig. 2A-B, Supplementary Fig. 4). The new full-length chimera H1H5_FL2 and H1H3_FL2 bound with an apparent K_D_ of 137 nM and approximately 368 nM to their respective strain-specific RBS-targeting mAb (Fig. 2C). While mAb F045 was reported to have cross reactivity to some H1 strains, we observed weak binding which could not be fit (Supplementary Fig. 5), therefore binding to H1H3_FL2 increased over H1SI06 is the result of the designed H3 RBS chimerism^7^ (Fig. 2D). The affinity of H1H3_FL2 to the H3-RBS-specific antibody did not vastly improve, yet both H1H5_FL2 and H1H3_FL2 are within 10-fold difference in affinity to the subtype-specific RBS antibody compared to their respective wildtype. Structural differences in the chimera RBS compared to the respective wildtype RBS, particularly at the 130-loop, may contribute to the difference in affinity. The RBS is a complex motif likely influenced by core residues in the HA head and replacement of only surface residues limits the ability to fully recapitulate the RBS structure, particularly of cross-group chimera. Nonetheless, the FL2 designs are stable in solution with a mixed alpha helix beta sheet profile by CD and are thermostable in contrast to the H1H3_Head design (Supplementary Fig. 6). These results highlight the benefit of using full length trimers to present increasingly radical RBS replacements.

**Figure 2:**
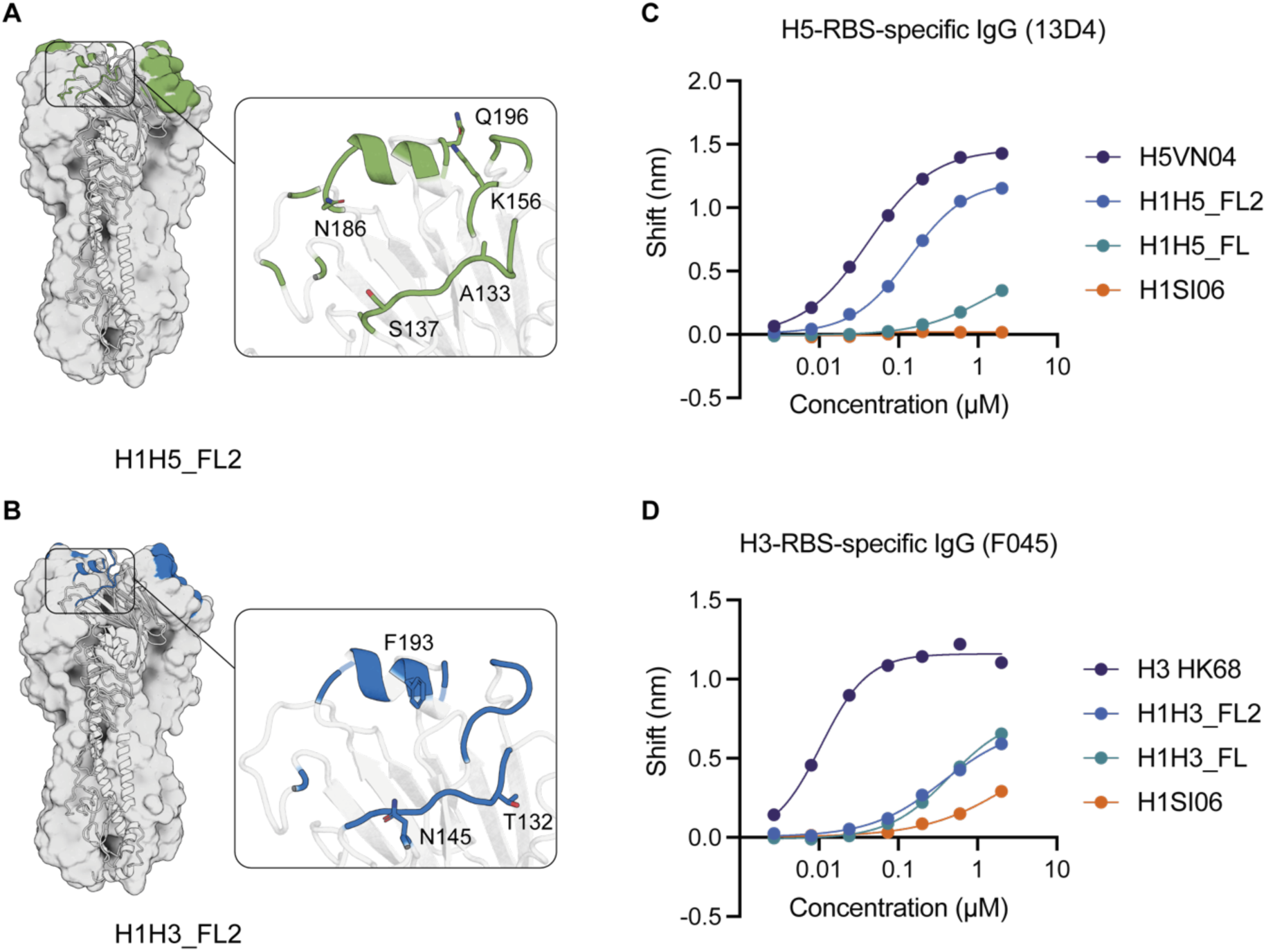
Full length RBS chimera bind subtype-specific antibodies. A-B) Structure of the H1SI06 hemagglutinin RBS epitope highlighting additional residues included in the full-length chimera. Transplanted residues by H5VN04 and H3HK68 are highlighted in green and blue, respectively. Left panel shows zoom in on RBS highlighting major contact residues added in H1H5_FL2 and H1H3_FL2 C-D) BLI affinity measurements for H5VN04, H1H5_FL, H1H5_FL2, and H1SI06 against an H5 RBS-subtype-specific antibody 13D4 IgG and H3HK68, H1H3_FL, H1H3_FL2, and H1SI06 against an H3 RBS-subtype-specific antibody F045-92 IgG, respectively.

### Chimeras provide heterotrimeric compatibility of cross-group RBS

The homotrimeric chimera presents cross-subtype diversity at the RBS, yet the remaining head surface, particularly the trimerization interface, is conserved. Leveraging this property, we hypothesised that these chimeras would facilitate the formation of heterotrimeric constructs presenting cross-group RBS. While intra-group heterotrimerisation is possible via linked tandem heterotrimers^15^, the diversity between group 1 and group 2 sequences can complicate incorporation into heterotrimeric constructs as a result of sequence variability at the trimerization interfaces. Our cross-group RBS chimera presents the conserved H1SI06 trimerization interface thereby facilitating heterotrimeric incorporation of our chimeric full-length HA from any cross-group RBS replacement, in which each of the three monomers represents a unique RBS (H1, H5, H3) H1_H1H5_H1H3 (Fig. 3A). Using tandem linkers^15^,we designed a H1_H1H5_H1H3 chimeric heterotrimer which shows a similar CD profile to full length H1SI06 WT and similar mass to a H1SI06 WT construct by mass photometry supporting the formation of trimer (Supplementary Fig. 7). The H1_H1H5_H1H3 heterotrimer binds an H1 RBS-specific antibody (CH65)^19^ with low nanomolar affinity (steady state apparent K_D_ >160 nM). The heterotrimer additionally binds a H5 RBS specific antibody (13D4)^18^ with nanomolar affinity and to an H3-specific antibody (F045-92 IgG)^7^ with micromolar affinity (steady state apparent >200 nM and >1 μM, respectively) (Fig. 3B-D). The decrease in affinities from the chimeric subunits may be the result of the instability in solution of the fused heterotrimer as opposed to using an exogenous trimerization domain resulting in instability of the chimeric head. The heterotrimer does show a mixed secondary structure profile by CD yet the linear melting curve may be indicative of transient stability in solution (Supplementary Fig. 7). Incorporation of an exogenous trimerization domain in the genetically fused construct may improve stability of the heterotrimer and improve presentation of the chimera subunits. Taken together, these results indicate that heterotrimeric assembly of RBS chimera allows for modularity in the presentation of cross-group RBS on one immunogen.

**Figure 3:**
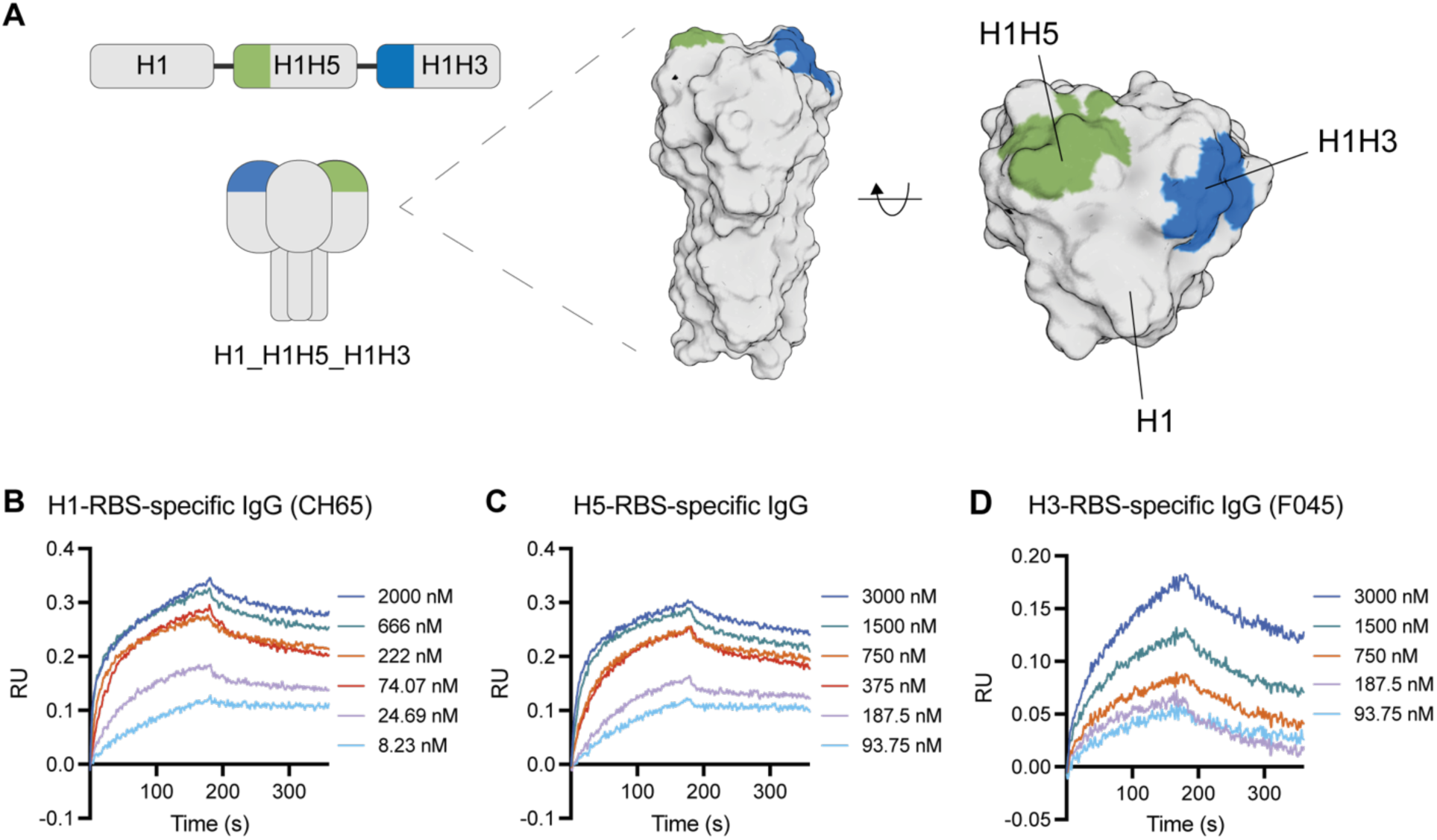
Heterotrimeric hemagglutinin chimera display cross-subtype and cross-group antibody binding. A) Linear construct of H1_H1H5_H1H3 chimeric heterotrimer and schematic. Left panel shows AlphaFold3 prediction of trimeric hemagglutinin consisting of the H1SI06 (grey), H1H5_FL2 (green), and H1H3_FL2 (blue). B-D) BLI affinity measurements against H1, H5, and H3 RBS-specific antibodies CH65 IgG, 13D4 IgG and F045-92 IgG, respectively.

### Homotrimeric chimera elicits cross-group reactivity

Given the ability of the chimeric heterotrimer to engage with three subtype-RBS-specific antibodies, we further assess the chimeric heterotrimer for the ability to elicit antibodies to each replaced subtype. We immunised 6 groups of mice (n=5) with a homologous boost scheme of H1SI06 WT, H5VN04 WT, H3HK68 WT, H1H5_FL2, H1H3_FL2, or H1_H1H5_H1H3 chimeric heterotrimer (Fig. 4A) and assessed reactivity to the subtype HA head domains. Both chimeric homotrimers (H1H5_FL2 and H1H3_FL2), as well as the H1_H1H5_H1H3 heterotrimer, show high serum IgG titers against H1 head domain (Fig. 4B). There no improvement of the serum IgG titers against H5 induced by H1H5_FL2 or H1_H1H5_H1H3 heterotrimer compared to base H1SI06, likely attributed to the existing cross-reactivity between H1 and H5 as closely related subtypes (Fig. 4C). On the other hand, the H1H3_FL2 chimera showed a significant improvement in H3 binding titers compared to H1SI06 WT (Fig. 4D). However, the H1_H1H5_H1H3 heterotrimer showed similar reactivity against the H3 head domain as H1SI06 WT did (Fig. 4C-4D), indicating that a large proportion of the antibody response targets the conserved H1SI06 base.

**Figure 4:**
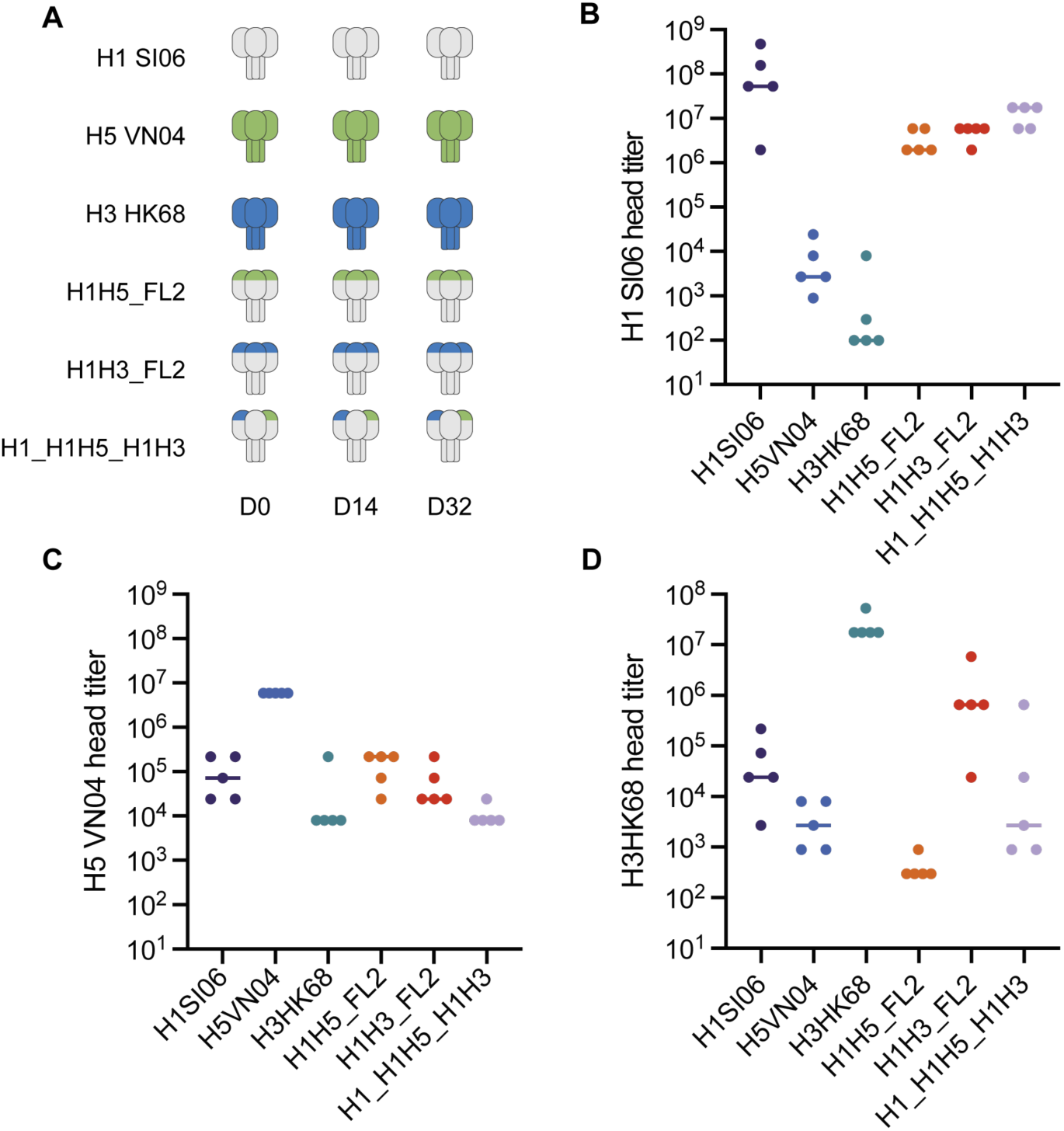
RBS chimera elicit cross-reactive antibodies. A) 6 groups of mice (n=5) were immunised three times in a homologous boost scheme with H1SI06, H5VN04, H3HK69, H1H5_FL2, H1H3_FL2, or H1_H1H5_H1H3 heterotrimer. B-D) Binding titers of the mouse serum IgG collected at day 56 post-prime against H1, H5 or H3 head domains determined by enzyme-linked immunosorbent assay (ELISA). The median values are shown as lines.

Given the cross-reactivity of the chimeric immunogens, we assessed the neutralisation potency of the immune sera. The elicited sera from all novel immunogens H1H5_FL2, H1H3_FL2, and H1_H1H5_H1H3 heterotrimer were able to neutralise a pre-pandemic H1N1 virus (A/Brisbane/59/2007) (Fig. 5A). Br07 was used as the Br07 HA possesses 98.3% sequence identity to the HA SI06. In addition, the sera from H1H5_FL2 showed low neutralising activity against H5N1 (NIBRG-14, carrying HA derived from A/Vietnam/1194/2004), yet higher than the H1SI06 sera (Fig. 5B). Likely the specific reactivity to H5 observed in ELISA by H1H5_FL2 was the contribution of the natural cross-reactivity of H1 and H5. However, increased neutralisation of H5N1 virus by H1H5_FL2 compared to H1SI06 corroborates that the antibodies elicited by H1H5_FL2 are specific to the H5 RBS beyond H1 cross-reactivity. Notably, the sera elicited by the chimeric immunogen H1H3_FL2 was able to neutralise H3N2 virus (A/X-31 vaccine strain) although less potently than the sera from homologous H3HK68 (Fig. 5C). The sera from the H1_H1H5_H1H3 heterotrimer neutralised H1N1 with a similar potency as H1SI06 WT immune sera, yet neutralisation of H5N1 or H3N2 was not detected at the tested serum dilutions (Fig. 5A-C). Taken together, these data support the capacity of chimeric immunogens to elicit a cross-group polyclonal response against desired strains as a single trimer immunogen but when presented as a heterotrimer common antigenic sites are dominant in the immunogenic response.

**Figure 5:**
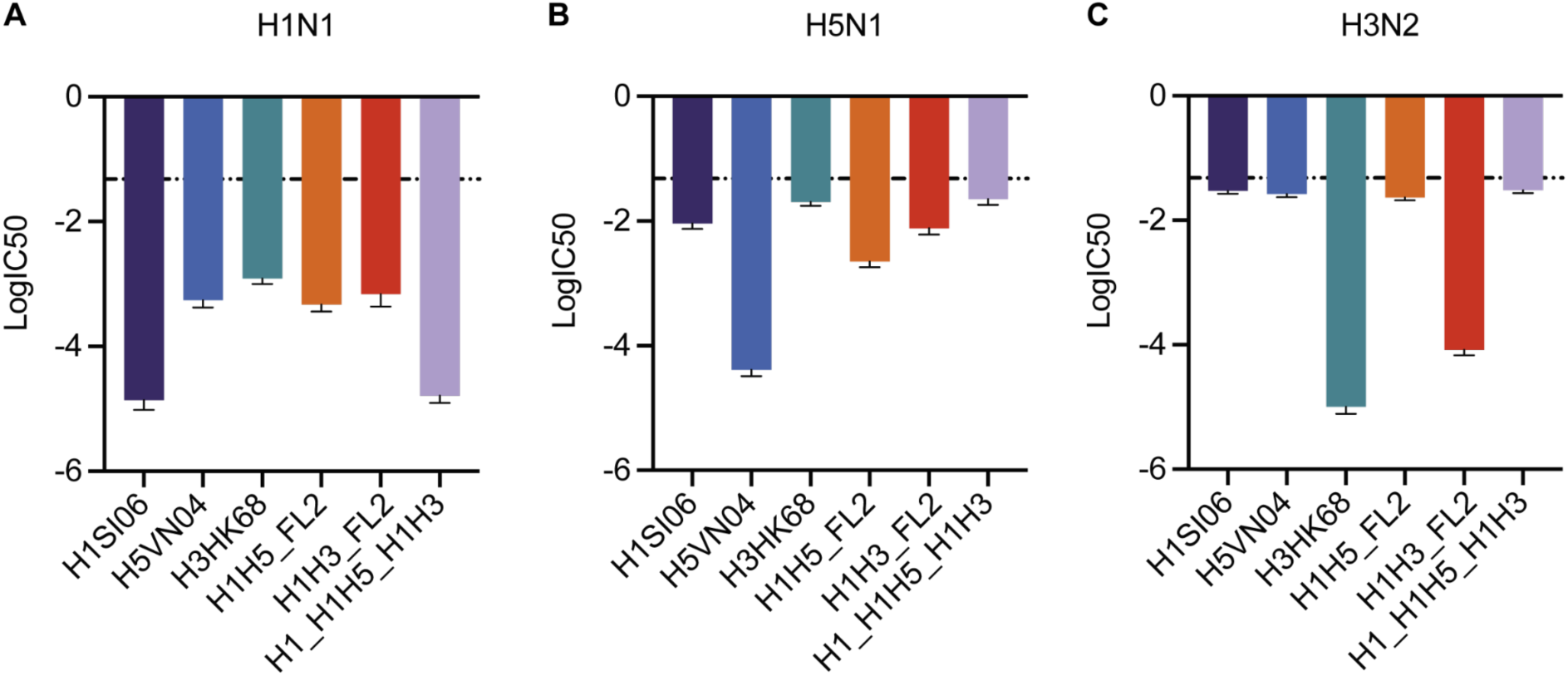
Cross-group neutralization elicited by RBS chimera. Neutralising activity of mouse immune sera raised against full length hemagglutinin WT, chimera, or heterotrimer against A) H1N1 (A/Brisbane/59/2007) pre-pandemic strain, B) H5N1 (NIBRG-14, carrying HA derived from A/Vietnam/1194/2004), C) H3N2 (A/X-31). The logarithm of IC_50_ values with standard error of the mean (s.e.m.) are shown. The threshold line represents day 0.

## Discussion

Few sites on the HA head are conserved in sequence between subtypes, emphasised by the relatively rare neutralising head antibodies that are cross-reactive to different subtypes. Yet, these major antigenic sites are well defined for group 1 and group 2 subtypes and are strikingly similar in structure. We leverage the definition of these sites to replace the major segments of the RBS antigenic site across subtypes. The RBS chimera represents a mid-strain sequence consisting of elements from both strains in the HA head. These chimeric head domains would likely not emerge in nature as their vulnerability to H1 and H5 specific antibodies could present a disadvantage to viral fitness. However, leveraging this property as an immunogen may be able to elicit antibody responses bridging two HA subtypes. In naive mice, our H1H3 chimeric immunogen was able to induce immune serum that could potently neutralise not only H1N1, but H3N2 virus. Currently, the RBS chimera only presents two strains of interest which could be used for probing immunogenicity of specific strains. Notably, the strains used in this study share moderate sequence identity to currently circulating strains obtained through GISAID^24–26^. Compared to 2024-2025 recommended vaccine strains, H1SI06 shares 72.7% and 72.4% sequence identity to A/Wisconsin/67/2022 (H1N1)pdm09-like virus and A/Victoria/4897/2022 (H1N1)pdm09-like virus respectively, while H3HK68 shares 83% sequence identity to both A/Thailand/8/2022 (H3N2)-like virus and A/Massachusetts/18/2022 (H3N2)-like virus. H5VN04, similarly, shares between 79.7% and 88.7% sequence identity to 2024 human-isolated H5 strains. The modularity presented by the RBS chimera allows for a rapid and flexible exchange of sequences and would be readily able to incorporate more recent circulating strains and could additionally incorporate consensus sequence strategies such as COBRA to improve the breadth of the exchanged RBS.

Current influenza vaccines incorporate HA of circulating strains as a cocktail vaccine. Here we propose a chimeric heterotrimer presenting three HA subtypes of pandemic interest as a single immunogen. The conservation of the HA head, apart from the major RBS antigenic site, encourages modularity of the chimeric heterotrimer without affecting heterotrimerization interfaces thereby allowing phylogenetically distinct strains to exist in a single trimer. While H1_H1H5_H1H3 heterotrimer showed low affinity to an H1-, H5- and H3-specific antibody, reduced from homotrimeric chimera, contribution from the H1 RBS to the binding of the H3- specific mAb F045-92 should be deduced with a H1 RBS mutant given the low cross-reactivity of the mAb to H1SI06. The H1_H1H5_H1H3 heterotrimer potently neutralised H1 but neutralisation was not detected by H5 or H3 head domain. Likely, the loss of avidity effects rendered the heterotrimer unable to engage with B-cell precursors targeting the RBS. Rather, the vast majority of antibodies were focused to the H1 conserved regions between the monomers and a small proportion targeted the H5 RBS or H3 RBS. For a heterotrimeric construct to support vaccine breadth by eliciting a polyclonal response to heterosubtypes, likely a larger sequence exchange will be required. Similar effects were observed in a study of heterotrimeric HIV constructs which benefited from improved neutralisation potency compared to homotrimeric counterparts, yet did not improve breadth^27^. While we do not observe improved breadth with homotypic immunisation with H1_H1H5_H1H3 heterotrimer, such an immunogen may present a pathway to target antibody evolution and may be of use in heterotypic immunisation for germline-targeting and polishing vaccination strategies^28^. Further study of the antibody repertoire elicited may elucidate the benefit of the proximity of cross-strain RBS domains of such cross-group heterotrimers.

Here we demonstrate that a single immunogen presenting at least one major antigenic site from two phylogenetically diverse strains effectively produces a neutralising response in mice. Cross-group responses are critical in the progress of a broadly protective influenza vaccine. More broadly, the chimeric approach could be applied to viruses with large mutagenicity potential yet conserved structure, including the spike protein of coronaviruses.

## Acknowledgements

We gratefully acknowledge all data contributors, i.e., the Authors and their Originating laboratories responsible for obtaining the specimens, and their Submitting laboratories for generating the genetic sequence and metadata and sharing via the GISAID Initiative, on which this research is based. We thank M. Pacesa for helpful discussions about the manuscript. We thank several EPFL facilities: PTPSP (F. Pojer, Y. Doo, K. Lau, L. Durrer, and S. Quinche) for protein expression, biochemical instrument support and the phenogenomics centre for support with mouse experiments (G. Diaceri). We additionally thank J. Williams for data transfer support.

This research used resources of the Advanced Photon Source; a U.S. Department of Energy (DOE) Office of Science User Facility operated for the DOE Office of Science by Argonne National Laboratory under Contract No. DE-AC02-06CH11357. Data were collected at Southeast Regional Collaborative Access Team (SER-CAT) 22-ID beamline at the Advanced Photon Source, Argonne National Laboratory. SER-CAT is supported by its member institutions (see www.ser-cat.org/members.html), and equipment grants (S10_RR25528 and S10_RR028976) from the National Institutes of Health. Funding: This work was sponsored by GlaxoSmithKline Biologicals SA, and was supported by the Swiss National Science Foundation: 310030_197724, by the ENDFLU project that has received funding from the European Union’s Horizon 2020 research and innovation program under Grant agreement No. 874650. The funders had no role in the preparation of the manuscript.

## Author Contribution

K.M.C. and B.E.C. conceived the work and designed the experiments. K.M.C. performed the chimera design. K.M.C., A.R., S.W., and S.X. performed the experimental characterizations. N.S., W.H., C.M. and V.V. contributed to structural studies and solved the crystal structures. K.M.C., S.G. and J.S. contributed to the design and planning of animal studies, and sera analysis. X.S. and H.C. designed and performed the neutralisation studies. K.M.C. and B.E.C. wrote and edited the manuscript with input from all authors.

## Data Availability

Structures are accessible in the Protein Data Bank under PDB ID 9MER (H1H5:FluA20 complex) and 9MEV (H1H3:FluA20 complex). Plasmid sequences are available in Supplementary Table 1. The plasmids of the designed proteins are available from the authors under a material transfer agreement with the Ecole Polytechnique Fédérale de Lausanne (EPFL). Accession identifiers for influenza sequences accessed through GISAID are summarized in Supplementary Table 3.

## Materials and Methods

### HA RBS chimera design, and structure and sequence analysis

Design of HA head domain chimeras were computationally generated using a structurally-guided segment replacement approach. The H1 strain (A/Solomon Islands/3/2006(H1N1)) (SI06) was used as the base construct. Rosetta LayerDesign^29,30^ was used to computationally identify core, surface and boundary residues based on solvent exposure which defined residues available for graft replacement on SI06 (PDB ID: 6OC3). Grafted strains for H5 and H3 were selected based the availability of a crystal structure, VN04 (A/Vietnam/1194/2004(H5N1)) (PDB ID: 4BGW) and (A/Hong Kong/1/1968(H3N2)) (HK68) (PDB ID: 6OCB) respectively. Vaccine strain sequences and human-isolated 2025 H5N1 seequences for comparison were obtained using GISAID accession codes listed in Supplementary Table 3.

The structures were aligned using Pymol Super^31^ to obtain a sequence independent Cα-based alignment. Consurf was used to determine the sequence variability between H1 and H5, or H1 and H3. A select dataset of sequences from 1960-2009 were selected from the Influenza virus database^32^ to form the multiple sequence alignment and consurf visualization. Replaced regions were defined as equivalent regions of known H1 epitopes with convergence on structural similarity to H5 or H3. If deviations were identified, the replacement segment was extended to regions of reconvergence. Rosetta-defined surface residues as well as solvent facing or inter-epitope facing boundary residues were selected for replacement. For FL2 designs, residues which made direct contact with the subtype-specific antibody were considered for subtype consensus residue identity rather than strain-specific residue identity. Residues which were not required for antibody binding were reverted to H1 native sequence to improve stability. The chimera designs were validated using Rosetta FastDesign to generate computational models and AlphaFold^33^ to validate the predicted structure.

### Multi-angle light scattering

The oligomeric state and molecular weight of the HA head chimeras were determined with size exclusion chromatography coupled to a multi-angle light scattering device (miniDAWN TREOS, Wyatt). Purified head domains were concentrated to 1 mg/ml in PBS (pH 7.4), and 100 μl of the sample was injected into a Superdex 75 300/10 GL column (Cytiva) with a flow rate of 0.5 ml/min, and UV280 and light scattering signals were recorded. Molecular weight was determined using the ASTRA software (version 6.1, Wyatt).

### Mass Photometry

Mass photometry experiments were conducted using a Refeyn TwoMP system (Refeyn Ltd., Oxford, UK) equipped with the AcquireMP and DiscoverMP software packages for data acquisition and analysis, respectively, with standard settings. For the experiments, microscope coverslips (Refeyn) of high precision were used on a one-time basis. To maintain the droplet shape of the sample, self-adhesive silicone culture wells (Grace Bio-Labs reusable CultureWell gaskets) were employed. For contrast-to-mass calibration, Bovine Serum Albumin Fraction V low Heavy Metals (Millipore) oligomers with molecular weights of 66, 132, 198, and 264 kDa were employed. Prior to the measurements, protein stocks were diluted in stock buffers containing 20 mM HEPES pH 7.5 and 150 mM NaCl. Specifically, 2 μL of the protein solution was combined with 18 μL of analysis buffer, resulting in a final drop volume of 20 μL with a concentration of approximately 1 μg/mL. Measurements were recorded for 60 seconds.

### Surface plasmon resonance

SPR measurements were performed on a Biacore 8K (GE Healthcare) in 10 mM HEPES pH 7.4, 150 mM NaCl, 3 mM EDTA, 0.005% v/v Surfactant P20 (GE Healthcare). Antibodies were immobilized on a CM5 chip (GE Healthcare # 29104988) by amine coupling. Approximately 500-1000 response units (RU) of protein were immobilized, and head chimera were injected as analyte in 2-fold serial dilutions. The flow rate was 30 μl/min for a contact time of 120 s followed by 600 s dissociation time. After each injection, the surface was regenerated using 0.1 M glycine at pH 3.0. Data were fitted using 1:1 Langmuir binding model within the Biacore 8K analysis software (GE Healthcare #29310604).

### Biolayer interferometry

BLI measurements were performed on a Gator bio BLI in 10 mM HEPES pH 7.4, 150 mM NaCl, 3 mM EDTA, 0.005% v/v Surfactant P20 (GE Healthcare). Full length hemagglutinin WT, chimera or heterotrimer was immobilized on anti-His probes (Gator Bio #160009) at 10 μg/mL for 180s. Antibody IgG was used as analyte at 6 μM or 2 μM in a 3-fold dilution series for an association time of 180s and dissociation of 180s. The probes were regenerated with 0.1 M glycine at pH 2.0. Data were fitted using 1:1 binding model within the GatorOne software (Gator bio).

### Circular dichroism

Circular dichroism spectra were measured using a Chirascan instrument in a 1-mm path-length cuvette. The protein samples were prepared in 10 mM sodium phosphate buffer at a protein concentration of ∼30 μM. Wavelengths between 195 nm and 250 nm were recorded with a scanning speed of 20 nm min−1 and a response time of 0.125 secs. All spectra were averaged two times and corrected for buffer absorption. Temperature ramping melts were performed from 20 to 90 °C with an increment of 2 °C/min or 5 °C/min. Thermal denaturation curves were determined by the change of ellipticity minimum at 220 nm.

### Hemagglutinin protein expression and purification

The full-length hemagglutinin trimers were cloned into pVRC vector (REF) and head constructs were cloned into pHLsec vector. The head domain plasmids and the wild-type full-length strain plasmid was transfected in HEK293-F cells while the full-length chimeric strains were transfected in ExpiCHO cells. Supernatants were harvested after 7 days and purified by Ni-NTA affinity chromatography eluting using 10 mM Tris, 500 mM NaCl, and 300 mM Imidazole (pH 7.5). Final purification by size exclusion chromatography in PBS (pH 7.4) on a Superdex 200 Increase 10/300 GL column (GE Healthcare).

### Antibody IgG and Fab protein expression and purification

Heavy and light chain DNA sequences of antibody fragments (Fab) were purchased from Twist Bioscience and cloned individually into the pHLsec mammalian expression vector (Addgene, #99845) using Gibson assembly. The heavy chain sequence was additionally cloned into pHLsec-Fc containing the human C_H_2 and C_H_3 region. HEK-293T cells were transfected with a 1:3 ratio of heavy to light chain. Supernatants were collected after 6 days and purified using a 5-ml HiTrap Protein A HP column (GE Healthcare) for IgG expression and 5-ml kappa-select column (GE Healthcare) for Fab purification. IgG or Fabs were eluted with 0.1 M glycine buffer (pH 2.7), immediately neutralized by 1 M Tris buffer (pH 9), and further purified by size exclusion chromatography on a Superdex 200 Increase 10/300 GL column (GE Healthcare) in PBS.

### Mouse immunizations

All animal experiments were approved by the Vaud Veterinary Cantonal Authorities in accordance with Swiss regulations of animal welfare (VD3808). Female BALB/c mice at 6-weeks old were obtained from Janvier labs and acclimatised for one week. Immunogens were mixed with equal volumes of adjuvant (AS03-like, Invivogen) and incubated for 1 hour on ice. Mice were injected subcutaneously with 50 uL of vaccine formulated with 2 ug of immunogen. Immunizations were performed on days 0, 21, and 42. Blood samples were collected on days 0, 14, and 35. Mice were euthanized on day 56 and blood collection was performed by cardiac puncture.

### ELISA

Purified recombinant full-length or head hemagglutinin was coated on Ninety-six well plates (Nunc MediSorp, Thermo Scientific) overnight at 4 C with 0.5 ug/mL in coating buffer (phosphate buffered saline (PBS) pH 7.4). Plates were washed three times at each step with wash buffer (PBS + 0.05% Tween-20 (PBS-T)) Wells were blocked with blocking buffer (PBS-T with 5% skim milk (Sigma)) for 1h at room temperature. Three-fold serial dilutions were prepared in assay buffer (PBS-T + 1 % Bovine Serum Albumin) and were added to the plates and incubated at room temperature for 2 hours. Secondary incubation with anti-mouse IgG (abcam, #99617) HRP-conjugated secondary antibody diluted 1:1,500 was added and incubated for 1 hour at room temperature. Plates were developed by adding 100 uL TMB substrate (Thermo Scientific). The reaction was stopped using and equal volume of 0.5 M HCl. The absorbance at 450nm was measured on a Tecan Safire 2 plate reader. The titer was determined as the reciprocal of the serum dilution which resulted in a signal two-fold above background.

### Microneutralization (MN) Assay

The microneutralization assay was performed on Madin-Darby canine kidney (MDCK) cells. A total of 20,000 MDCK cells were seeded per well in 96-well plates 24 hours before virus inoculation. Mouse serum samples from day 0 and day 56 after immunisation were pooled per immunisation group. The serum samples were heat treated at 56°C for 30 minutes to inactivate complement.

Human monoclonal antibody CR9114 (hmAb)^34^ and related convalescent mouse serum were used as positive control. The mouse serum collected on day 0 was used as a negative control. Virus stocks of H1N1 (A/Brisbane/59/2007), H5N1 (NIBRG-14, recombinant virus with HA and NA from A/Vietnam/1194/2004) and H3N2 (A/Hong Kong/1/1968) were used after optimising the inoculum titers for optimal infection conditions. The maintenance DMEM medium (Gibco, high glucose) contains NEAA Culture Supplement, ampicillin (100μg/ml) and 1ug/ml TPCK-treated trypsin (Sigma).

Serum samples and CR9114 were 3-fold diluted in 96-well plates with the maintenance medium. The starting concentration of CR9114 was 20µg/ml and that of the mouse serum was 1/10 [HC1] dilution. Virus samples were combined with serial dilutions of serum/mAb and the mixtures were incubated at 37°C for 30 min. After incubation, 100ul of virus/antibody mix were transferred to MDCK cells. Forty-eight hours later, the cell medium was removed and plates were washed with PBS. Cells were then fixed with 4% paraformaldehyde (in PBS). After two washing steps with PBS, the cells were permeabilized by adding PBS with 0,5% Triton X-100, followed by incubation for 10 min at room temperature, after which the wells were blocked by the addition of PBS with 0,05% Tween20 and 3% milk powder, followed by incubation for 1 hour at room temperature. Then cells were stained with Polyclonal Anti-Influenza Virus RNP (BEI Resources NR3133) diluted 1/2000 in PBS with 0,05% Tween20 for 1 hour at room temperature. After 3 washes with PBS with 0,05% Tween20, the secondary HRP-antibody (Donkey anti-goat-HRP, Ab97110) diluted 1/5000 in PBS with 0,05% Tween20 was added and incubated for 1 hour at room temperature. Then plates were washed three times with PBS and developed with TMB substrate (BD OptEIA™ TMB Substrate Reagent Set). The absorbance at 450 nm was determined by iMark™ Microplate Absorbance Reader (BIO-RAD).

The absorbance (450 nm) data were analyzed using GraphPad Prism (version 9.5.1). The lowest value in each subcolumn was defined as 0% and the highest values were defined as 100% for the normalisation. Based on the normalised data, IC_50_ of serum samples were calculated with Nonlinear Regression using model log(inhibitor) vs. normalised response. LogIC_50_ values of the serum samples from day 56 were plotted according to the different virus strains. The baseline was set as the average value of the day 0 serum samples.

### Expression and purification of H1H5 HA, H1H3 HA, and FluA20 Fab proteins for crystal structure determination

FluA20 Fab, H1H5 and H1H3 Hemagglutinin chimeric antigens were expressed in Expi293F GnTI- HEK (Gibco catalog # A39240) mammalian cells by transient transfection separately, following manufacturer’s protocol. In brief, Expi293F GnTI- HEK cells, derived from Expi293F cells but engineered to lack N-acetylglucosaminyl transferase for uniform glycosylation patterns, were maintained in Expi293 Expression media (Gibco catalog # A14351-01) and grown at 37°C, 8% CO_2_, 80% humidity at 110 RPM. The day before transfection, cells were seeded at 2.4 × 10^6^ cells/mL after a complete media exchange. On the day of transfection, the cell density and viability were measured by trypan blue exclusion on a ViCell XR cell viability analyzer. The culture was adjusted to 3.6–3.8 × 10^6^ cells/mL by medium feed or culture split.

DNA plasmids were diluted in prewarmed Opti-MEM media (Gibco catalog # 11058-021) in a volume that was 5% of the transfection volume. 1 ug of DNA was transfected for every mL of transfection culture. ExpiFectamine 293 reagent (from Gibco ExpiFectamine 293 Transfection Kit, 1L, catalog # A14524) was diluted into a volume of prewarmed Opti-MEM media corresponding to 5% of transfection volume of prewarmed Opti-MEM.

Five minutes after the dilution of ExpiFectamine reagent in Opti-MEM, the diluted DNA solution was added to the diluted ExpiFectamine to form the DNA:ExpiFectamine complex for transfection. Twenty minutes after the addition of DNA to the ExpiFectamine reagent, the complex was added to the cell culture. Sixteen to twenty-two hours after the addition of diluted DNA to the cells, Enhancer 1 and 2 from the same kit were added to cell culture. Four to six days post transfection, cell count, and cell viability were measured with ViCell XR cell Viability Analyzer. When cell viability reached below 80%, cell media supernatants were harvested. The culture was centrifuged at 4200 rpm for 50 min at 10 °C and the supernatant was poured off into a clean 1L storage bottle. The supernatant was weighed to determine harvest volume and stored at 4 °C or −20 °C until purification.

The culture supernatant was loaded onto a pre-equilibrated HisTrap Excel column (Cytiva Life Sciences) at 5 ml/min using an AKTA Avant with 5 CV of Buffer A, (10 mM HEPES pH 8.0, 150 mM NaCl). The column was washed with another 5 CV of Buffer A, prior to elution. A combination of step-wise and linear gradient was performed with Buffer B, (10 mM HEPES pH 8.0, 150 mM NaCl, 500 mM Imidazole). Protein eluted at ∼100 mM imidazole. Fractions containing protein were pooled and concentrated prior to Size-Exclusion Chromatograph (SEC). SEC was performed using a Superdex 200 Increase 10/300 GL (Cytiva Life Sciences) with 150 mM sodium chloride, 10 mM HEPES, pH 7.5 at 0.5 ml/min. Protein purity and yield was determined by SDS-PAGE and concentration measured on NanoDrop 8000 using the respective extinction coefficients. Protein was aliquoted and stored at −80°C until further use.

### H1H5:FluA20 and H1H3:FluA20 complex formation and crystallization

Either H1H5 or H1H3 antigens were complexed with FluA20 in a 1:1.2 molar ratio, respectively and equilibrated at 4°C overnight for complex formation. Complexes were then subject to Size-Exclusion Chromatography on a HiLoad Superdex 200 Increase 16/600 GL (Cytiva Life Sciences) pre-equilibrated with 10 mM HEPES pH 7.5, 150 mM NaCl at 1 ml/min and eluted in the same buffer, to separate complex from antigen and fab. Peak fractions were run on SDS-PAGE and fractions containing complex were pooled and concentrated to 10 mg/ml as determined by their extinction coefficients on a NanoDrop 8000.

Protein complexes were screened for crystals using several commercially-available crystallization screening kits, (JCSG-Plus, Peg/Ion, Index, SG1) using a sitting drop vapor diffusion method, testing both 1:1 and 2:1 ratio of protein to reservoir solution with a starting concentration of 10 mg/ml of complex and a reservoir volume of 50 uL. Crystal trays were allowed to incubate at 20°C and visualized every couple of days for crystals in a RockImager. Diffraction-quality crystals appeared between 10-15 days and were assessed to be protein crystals by UV.

### Crystal structure determination

Diffraction quality crystals of H1H5:FluA20 and H1H3:FluA20 complexes were grown in 0.2 M NH4NO3, 20% w/v PEG 3350 and 0.1 M PCB pH 6, 25% w/v PEG 1500, respectively. Crystals were cryo-preserved with 10% ethylene glycol, before being sent off for data collection. Data was collected at the Southeast Regional Collaborative Access Team (SER-CAT) 22-ID beamline at the Advanced Photon Source, Argonne National Laboratory. The crystals were processed and diffracted to 2.00 and 2.06 Å, respectively. The crystals belonged to space group P 43 21 2 and P 1 21 1, respectively and contained a single copy of the antigen:fab complex in the asymmetric unit. Molecular replacement using Phenix PHASER program was used to resolve phases using PDB ID: 4BGW and 6OCB for H1H5 and H1H3 HA, respectively. For FluA20, PBD ID 6OBZ was used for molecular replacement^35^. Iterative rounds of refinement was performed using Coot^36^, and Phenix REFINE. Validation was performed using MOLPROBITY. The final refinement statistics are summarized in Supplementary Table 2.

## References

1. Grech, V. & Borg, M. Influenza vaccination in the COVID-19 era. Early Human Development 148, 105116 (2020).

2. Boyoglu-Barnum, S. et al. Quadrivalent influenza nanoparticle vaccines induce broad protection. Nature 592, 623–628 (2021).

3. Yamayoshi, S. & Kawaoka, Y. Current and future influenza vaccines. Nat Med 25, 212–220 (2019).

4. Turbelin, C. et al. Age Distribution of Influenza Like Illness Cases during Post-Pandemic A(H3N2): Comparison with the Twelve Previous Seasons, in France. PLOS ONE 8, e65919 (2013).

5. Angeletti, D. & Yewdell, J. W. Understanding and Manipulating Viral Immunity: Antibody Immunodominance Enters Center Stage. Trends Immunol 39, 549–561 (2018).

6. Ekiert, D. C. et al. Cross-neutralization of influenza A viruses mediated by a single antibody loop. Nature 489, 526–532 (2012).

7. Lee, P. S. et al. Receptor mimicry by antibody F045–092 facilitates universal binding to the H3 subtype of influenza virus. Nat Commun 5, 3614 (2014).

8. Raymond, D. D. et al. Conserved epitope on influenza-virus hemagglutinin head defined by a vaccine-induced antibody. Proc. Natl. Acad. Sci. U.S.A. 115, 168–173 (2018).

9. Kanekiyo, M. et al. Mosaic nanoparticle display of diverse influenza virus hemagglutinins elicits broad B cell responses. Nature Immunology 20, 362–372 (2019).

10. Allen, J. D. & Ross, T. M. Bivalent H1 and H3 COBRA Recombinant Hemagglutinin Vaccines Elicit Seroprotective Antibodies against H1N1 and H3N2 Influenza Viruses from 2009 to 2019. Journal of Virology 96, e01652–21 (2022).

11. Sanchez, P. L., Andre, G., Antipov, A., Petrovsky, N. & Ross, T. M. Advax-SM^TM^-Adjuvanted COBRA (H1/H3) Hemagglutinin Influenza Vaccines. Vaccines 12, 455 (2024).

12. Cohen, A. A. et al. Construction, characterization, and immunization of nanoparticles that display a diverse array of influenza HA trimers. PLOS ONE 16, e0247963 (2021).

13. Giles, B. M. & Ross, T. M. A computationally optimized broadly reactive antigen (COBRA) based H5N1 VLP vaccine elicits broadly reactive antibodies in mice and ferrets. Vaccine 29, 3043–3054 (2011).

14. Schmidt, A. G. A modular platform to display multiple hemagglutinin subtypes on a single immunogen.

15. del Moral-Sánchez, I. et al. Triple tandem trimer immunogens for HIV-1 and influenza nucleic acid-based vaccines. npj Vaccines 9, 1–18 (2024).

16. Bajic, G. et al. Structure-Guided Molecular Grafting of a Complex Broadly Neutralizing Viral Epitope. ACS Infect. Dis. 6, 1182–1191 (2020).

17. Caradonna, T. M. et al. An epitope-enriched immunogen expands responses to a conserved viral site. Cell Reports 41, 111628 (2022).

18. Lin, Q. et al. Structural Basis for the Broad, Antibody-Mediated Neutralization of H5N1 Influenza Virus. Journal of Virology 92, 10.1128/jvi.00547-18 (2018).

19. Whittle, J. R. R. et al. Broadly neutralizing human antibody that recognizes the receptor-binding pocket of influenza virus hemagglutinin. Proceedings of the National Academy of Sciences 108, 14216–14221 (2011).

20. Bangaru, S. et al. A Site of Vulnerability on the Influenza Virus Hemagglutinin Head Domain Trimer Interface. Cell 177, 1136–1152.e18 (2019).

21. Ermler, M. E. et al. Chimeric Hemagglutinin Constructs Induce Broad Protection against Influenza B Virus Challenge in the Mouse Model. J Virol 91, e00286–17 (2017).

22. Nachbagauer, R. et al. A chimeric hemagglutinin-based universal influenza virus vaccine approach induces broad and long-lasting immunity in a randomized, placebo-controlled phase I trial. Nat Med 27, 106–114 (2021).

23. Hai, R. et al. Influenza Viruses Expressing Chimeric Hemagglutinins: Globular Head and Stalk Domains Derived from Different Subtypes. Journal of Virology 86, 5774–5781 (2012).

24. Khare, S. et al. GISAID’s Role in Pandemic Response. CCDCW 3, 1049–1051 (2021).

25. Elbe, S. & Buckland-Merrett, G. Data, disease and diplomacy: GISAID’s innovative contribution to global health. Global Challenges 1, 33–46 (2017).

26. Shu, Y. & McCauley, J. GISAID: Global initiative on sharing all influenza data – from vision to reality. Eurosurveillance 22, 30494 (2017).

27. Sellhorn, G. et al. Engineering, Expression, Purification, and Characterization of Stable Clade A/B Recombinant Soluble Heterotrimeric gp140 Proteins. Journal of Virology 86, 128–142 (2012).

28. Haynes, B. F., Kelsoe, G., Harrison, S. C. & Kepler, T. B. B-cell–lineage immunogen design in vaccine development with HIV-1 as a case study. Nat Biotechnol 30, 423–433 (2012).

29. Tyka, M. D. et al. Alternate states of proteins revealed by detailed energy landscape mapping. J Mol Biol 405, 607–618 (2011).

30. Fleishman, S. J. et al. Rosettascripts: A scripting language interface to the Rosetta Macromolecular modeling suite. PLoS ONE 6, 1–10 (2011).

31. PyMOL | pymol.org. https://www.pymol.org/.

32. Information, N. C. for B., Pike, U. S. N. L. of M. 8600 R., MD, B. & Usa, 20894. Influenza virus database - NCBI. https://www.ncbi.nlm.nih.gov/genomes/FLU/Database/nph-select.cgi?go=database.

33. Jumper, J. et al. Highly accurate protein structure prediction with AlphaFold. Nature 596, 583–589 (2021).

34. Beukenhorst, A. L. et al. The influenza hemagglutinin stem antibody CR9114: Evidence for a narrow evolutionary path towards universal protection. Front. Virol. 2, (2022).

35. Adams, P. D. et al. PHENIX: a comprehensive Python-based system for macromolecular structure solution. Acta Crystallogr D Biol Crystallogr 66, 213–221 (2010).

36. Emsley, P., Lohkamp, B., Scott, W. G. & Cowtan, K. Features and development of Coot. Acta Crystallogr D Biol Crystallogr 66, 486–501 (2010).

