## supplementary_materials for "Structure-based Design of Chimeric Influenza Hemagglutinins to Elicit Cross-group Immunity"

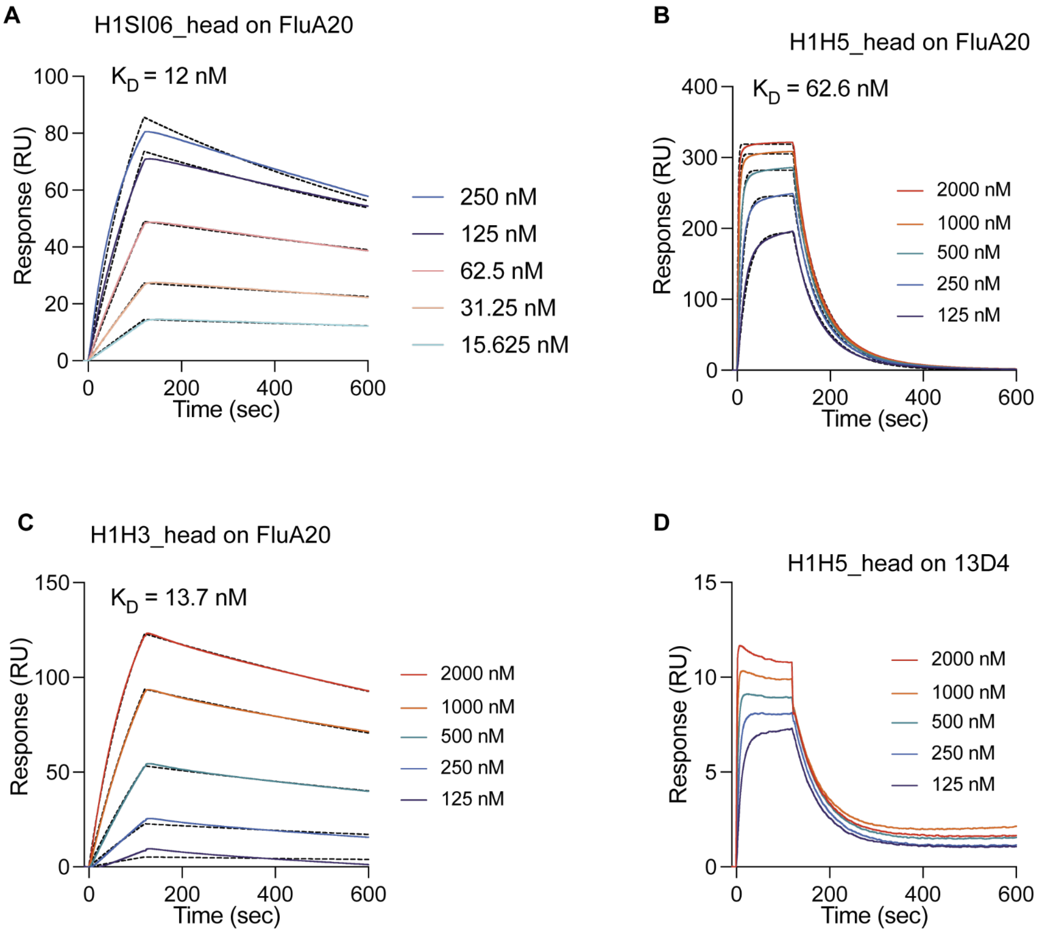


**Supplementary Figure 1: Biochemical characterization of HA head chimera**

A-C) SPR binding affinity of H1SI06_head,H1H5_head and H1H3_head to FluA20 IgG, respectively. D) SPR binding affinity of H1H5_head to H5-RBS-specific IgG 13D4. Curves could not be fit.

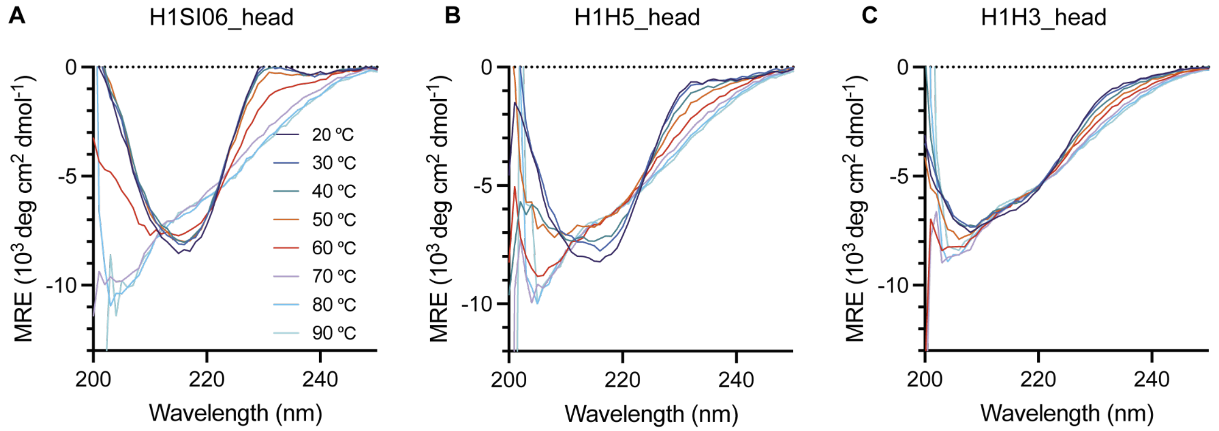


**Supplementary Figure 2: Circular dichroism of HA head chimera**

A-C) CD measured thermal stability of H1SI06_head, H1H5_head, and H1H3_head, respectively.

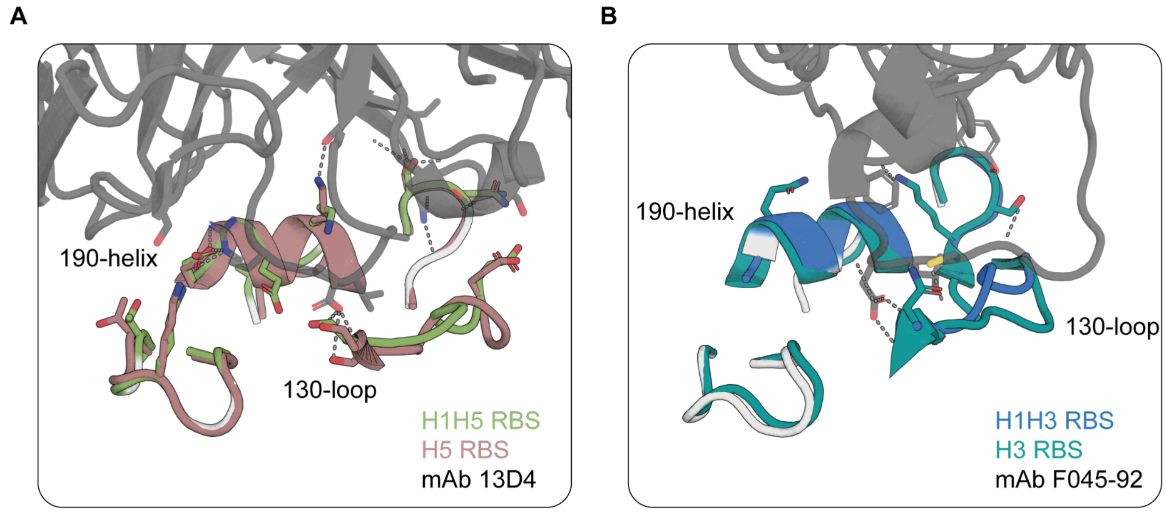


**Supplementary Figure 3: Antibody contact residues for RBS**

A) Crystal structure of the H1H5_head RBS (green) overlaid on H5 RBS (red) in complex with 13D4 Fab (grey). B) Crystal structure of the H1H3_head RBS (blue) overlaid on H3 RBS (teal) in complex with F045-92 Fab (grey).

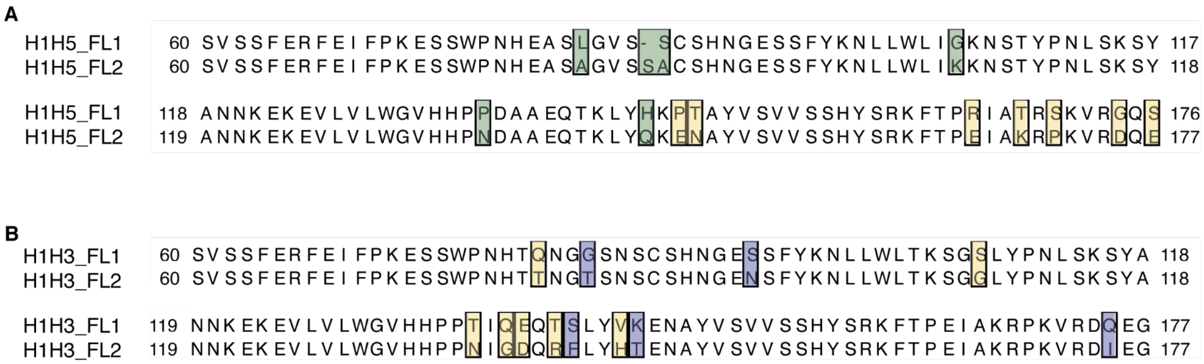


**Supplementary Figure 4: Full length chimera sequence alignment**

A) Sequence alignment of H1H5_FL1 to H1H5_FL2. Mutated residues to H5 sequences (green) are highlighted. B) Sequence alignment of H1H5_FL1 to H1H5_FL2. Mutated residues to H3 sequences (blue) are highlighted. Residues not directly contributing to RBS-antibody binding were reverted to H1SI06 identities (yellow).

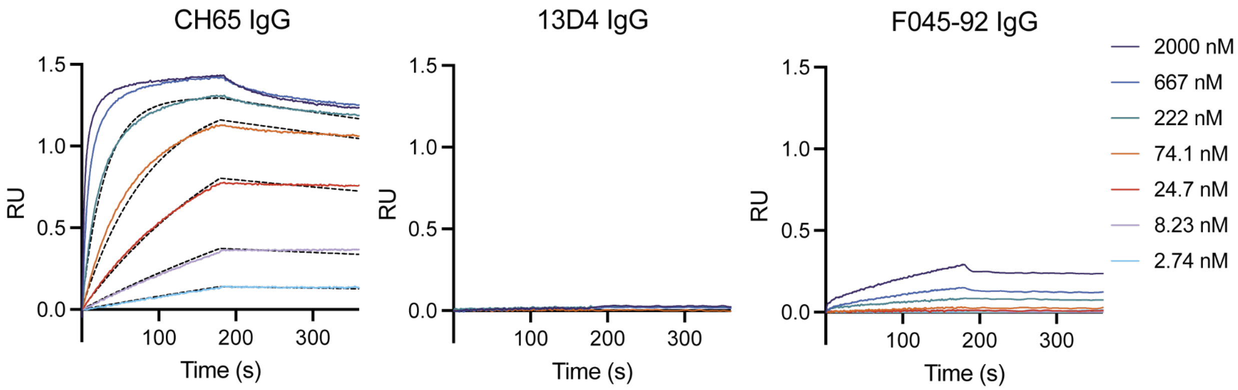


**Supplementary Figure 5: H1SI06 wild-type affinity to RBS antibodies**

A) BLI response of H1SI06_FL to CH65 IgG (H1-RBS binding), 13D4 IgG (H5-RBS binding), and F045 IgG (H3-RBS binding). F045-92 IgG kinetics could not be fit.

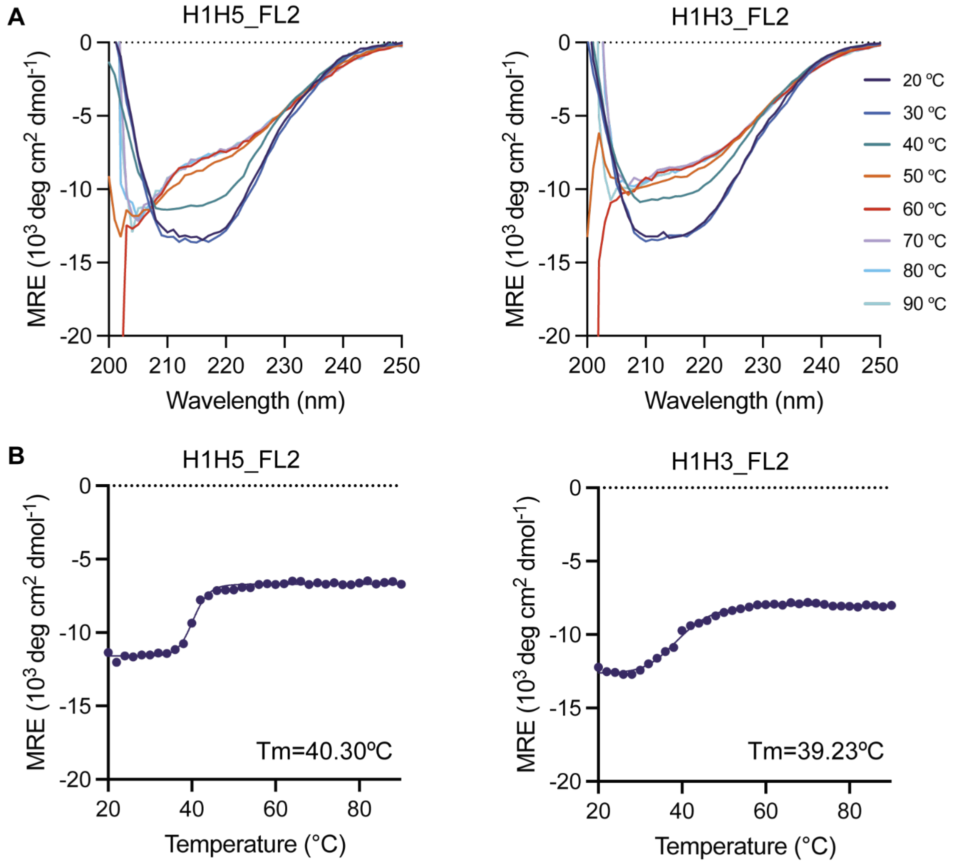


**Supplementary Figure 6: Biochemical characterization of HA full length chimera**

A) CD spectra of H1H5_FL2 and H1H5_FL2. B) Thermal stability of H1H5_FL2 and H1H5_FL2 with melting temperatures at 40.3 °C and 39.23 °C, respectively.

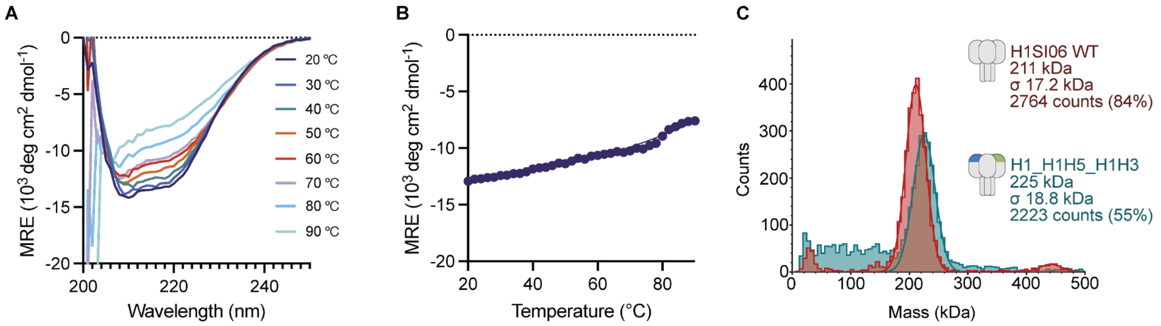


**Supplementary Figure 7: Structural analysis of H1_H1H5_H1H3 heterotrimer**

A) CD spectra of H1_H1H5_H1H3 chimeric heterotrimer. B) Thermal stability of H1_H1H5_H1H3 with a linear melting profile. C) Mass photometry comparing experimental masses of H1SI06 WT trimer and H1_H1H5_H1H3 heterotrimer showing apparent masses of 211 kDa and 225 kDa respectively.

**Supplementary Table 1: Crystal collection and refinement statistics**

| | H1H5_head in complex with FluA20 Fab | H1H3_head in complex with FluA20 |
|---|---|---|
| PDB ID: | 9MER | 9MEV |
| Wavelength | 1.0001 | 1.0000 |
| Resolution range | 43.1 - 2.0 (2.072 - 2.0) | 44.82 - 2.06 (2.09 - 2.06) |
| Space group | P 43 21 2 | P 1 21 1 |
| Unit cell | 79.533 79.533 256.452 90 90 90 | 50.641 93.465 86.434 90 93.482 90 |
| Total reflections | 455917 | 71760 |
| Unique reflections | 55274 (1587) | 45825 (1032) |
| Completeness (%) | 97.16 (78.83) | 92.14 (56.03) |
| Mean I/sigma(I) | 21.6 (3.1) | 17.9 (2.2) |
| Wilson B-factor | 23.83 | 24.96 |
| R-meas | 0.091(0.940) | 0.067 (0.521) |
| R-pim | 0.031 (0.374) | 0.033 (0.280) |
| CC1/2 | 0.996 (0.791) | 0.987 (0.811) |
| Reflections used in refinement | 55274 (1587) | 45825 (1032) |
| Reflections used for R-free | 2000 (57) | 1993 (48) |
| R-work | 0.2058 (0.2855) | 0.1809 (0.3106) |
| R-free | 0.2151 (0.2833) | 0.2091 (0.3444) |
| Number of non-hydrogen atoms | 5705 | 5728 |
| macromolecules | 5095 | 5073 |
| ligands | 130 | 66 |
| solvent | 480 | 589 |

| | | |
|---|---|---|
| Protein residues | 655 | 652 |
| RMS(bonds) | 0.014 | 0.011 |
| RMS(angles) | 1.65 | 1.29 |
| Ramachandran favored (%) | 96.61 | 96.89 |
| Ramachandran allowed (%) | 3.39 | 2.95 |
| Ramachandran outliers (%) | 0 | 0.16 |
| Rotamer outliers (%) | 0.17 | 0.34 |
| Clashscore | 12.54 | 5.72 |
| Average B-factor | 30.65 | 32.36 |
| macromolecules | 29.87 | 31.20 |
| ligands | 39.22 | 51.53 |
| solvent | 36.67 | 39.83 |

Statistics for the highest-resolution shell are shown in parentheses.

**Supplementary Table 2: Sequences of chimeric HA**

| Name | Sequence |
|---|---|
| H1H5_head | APLQLGNCSVAGWILGNPECELLISRESWS YIVEKPNPENGTCFPGHFADYEELREQLSS VSSFERFEIFPKESSWPNHEASLGVSSCSH NGESSFYKNLLWLIGKNSTYPNLSKSYANN KEKEVLVLWGVHHPPDAAEQTKLYHKPTAY VSVVSSHYSRKFTPRIATRSKVRGQSGRIN YYWTLLEPGDTIIFEANGNLIAPRYAFALSR GSGLVPRGSGHHHHHH |
| H1H3_head | APLQLGNCSVAGWILGNPECELLISRESWS YIVEKPNPENGTCYPGHFADYEELREQLSS VSSFERFEIFPKESSWPNHTQNGGSNSCS HNGESSFYKNLLWLTKSGSLYPNLSKSYAN NKEKEVLVLWGVHHPPTIQEQTSLYVKENA YVSVVSSHYSRKFTPEIAKRPKVRDQEGRI NYYWTLLEPGDTIIFEANGNLIAPRYAFALSR GSGLVPRGSGHHHHHH |

| | |
|---|---|
| H1H5_FL2 | DTVDTVLEKNVTVTHSVNLLEDKHNGKLCL<br>LKGIAPLQLGNCSVAGWILGNPECELLISRE<br>SWSYIVEKPNPENGTCFPGHFADYEELREQ<br>LSSVSSFERFEIFPKESSWPNHEASAGVSS<br>ACSHNGESSFYKNLLWLIKKNSTYPNLSKS<br>YANNKEKEVLVLWGVHHPNDAAEQTKLYQ<br>KENAYVSVVSSHYSRKFTPEIAKRPKVRDQ<br>EGRINYYWTLLEPGDTIIFEANGNLIAPRYAF<br>ALSRGFGSGIINSNAPMDECNTTCQTPEGAI<br>NTSLPFQNVHPITIGKCPKYVKSTKLRLATG<br>LRNVPSIQSRGLFGAIAGFIEGGWTGMVDG<br>WYGYHHQNEQGSGYAADLKSTQNAIDKITN<br>KVNSVIEKMNTQFTAVGKEFNHLEKRIENLN<br>KKVDDGFLDIWTYNAELLVLLENERTLDYHD<br>SNVKNLYEKVRNQLKNNAKEIGNGCFEFYH<br>KCDNTCMESVKNGTYDYPKYSEEAKLNRE<br>KIASGSGYIPEAPRDGQAYVRKDGEWVLLS<br>TFLGSHHHHHH |
| H1H3_FL2 | APLQLGNCSVAGWILGNPECELLISRESWS<br>YIVEKPNPENGTCYPGHFADYEELREQLSS<br>VSSFERFEIFPKESSWPNHTTNGTSNSCSH<br>NGENSFYKNLLWLTKSGGLYPNLSKSYANN<br>KEKEVLVLWGVHHPPNIGDQRFLYHTENAY<br>VSVVSSHYSRKFTPEIAKRPKVRDIEGRINY<br>YWTLLEPGDTIIFEANGNLIAPRYAFALSRGF<br>GSGIINSNAPMDECNTTCQTPEGAINTSLPF<br>QNVHPITIGKCPKYVKSTKLRLATGLRNVPSI<br>QSRGLFGAIAGFIEGGWTGMVDGWYGYHH<br>QNEQGSGYAADLKSTQNAIDKITNKVNSVIE<br>KMNTQFTAVGKEFNHLEKRIENLNKKVDDG<br>FLDIWTYNAELLVLLENERTLDYHDSNVKNL<br>YEKVRNQLKNNAKEIGNGCFEFYHKCDNTC<br>MESVKNGTYDYPKYSEEAKLNREKIDASGS<br>GYIPEAPRDGQAYVRKDGEWVLLSTFLGSH<br>HHHHH |
| H1_H1H5_H1H3 | DTLCIGYHANNSTDTVDTVLEKNVTVTHSV<br>NLLEDKHNGKLCLLKGIAPLQLGNCSVAGWI<br>LGNPECELLISRESWSYIVEKPNPENGTCYP<br>GHFADYEELREQLSSVSSFERFEIFPKESS<br>WPNHTTTGVSASCSHNGESSFYKNLLWLT<br>GKNGLYPNLSKSYANNKEKEVLVLWGVHH<br>PPNIGDQRALYHKENAYVSVVSSHYSRKFT<br>PEIAKRPKVRDQEGRINYYWTLLEPGDTIIFE<br>ANGNLIAPRYAFALSRGFGSGIINSNAPMDE<br>CNTTCQTPEGAINTSLPFQNVHPITIGKCPK<br>YVKSTKLRLATGLRNVPSIQSRGLFGAIAGFI<br>EGGWTGMVDGWYGYHHQNEQGSGYAAD<br>LKSTQNAIDKITNKVNSVIEKMNTQFTAVGK |

| | |
|---|---|
| | EFNHLEKRIENLNKKVDDGFLDIWTYNAELL<br>VLLENERTLDYHDSNVKNLYEKVRNQLKNN<br>AKEIGNGCFEFYHKCDNTCMESVKNGTYD<br>YPKYSEEAKLNREKIDGSGGSGGSGSGGS<br>GGSGGSGSGASDTLCIGYHANNSTDTVDT<br>VLEKNVTVTHSVNLLEDKHNGKLCLLKGIAP<br>LQLGNCSVAGWILGNPECELLISRESWSYIV<br>EKPNPENGTCFPGHFADYEELREQLSSVSS<br>FERFEIFPKESSWPNHEASAGVSSACSHNG<br>ESSFYKNLLWLIKKNSTYPNLSKSYANNKEK<br>EVLVLWGVHHPNDAAEQTKLYQKENAYVS<br>VVSSHYSRKFTPEIAKRPKVRDQEGRINYY<br>WTLLEPGDTIIFEANGNLIAPRYAFALSRGF<br>GSGIINSNAPMDECNTTCQTPEGAINTSLPF<br>QNVHPITIGKCPKYVKSTKLRLATGLRNVPSI<br>QSRGLFGAIAGFIEGGWTGMVDGWYGYHH<br>QNEQGSGYAADLKSTQNAIDKITNKVNSVIE<br>KMNTQFTAVGKEFNHLEKRIENLNKKVDDG<br>FLDIWTYNAELLVLLENERTLDYHDSNVKNL<br>YEKVRNQLKNNAKEIGNGCFEFYHKCDNTC<br>MESVKNGTYDYPKYSEEAKLNREKIGSGGS<br>GGSGSGGSGGSGGSGSGHMDTLCIGYHA<br>NNSTDTVDTVLEKNVTVTHSVNLLEDKHNG<br>KLCLLKGIAPLQLGNCSVAGWILGNPECELL<br>ISRESWSYIVEKPNPENGTCYPGHFADYEE<br>LREQLSSVSSFERFEIFPKESSWPNHTTNG<br>TSNSCSHNGENSFYKNLLWLTKSGGLYPNL<br>SKSYANNKEKEVLVLWGVHHPPNIGDQRFL<br>YHTENAYVSVVSSHYSRKFTPEIAKRPKVR<br>DIEGRINYYWTLLEPGDTIIFEANGNLIAPRY<br>AFALSRGFGSGIINSNAPMDECNTTCQTPE<br>GAINTSLPFQNVHPITIGKCPKYVKSTKLRLA<br>TGLRNVPSIQSRGLFGAIAGFIEGGWTGMV<br>DGWYGYHHQNEQGSGYAADLKSTQNAIDK<br>ITNKVNSVIEKMNTQFTAVGKEFNHLEKRIE<br>NLNKKVDDGFLDIWTYNAELLVLLENERTLD<br>YHDSNVKNLYEKVRNQLKNNAKEIGNGCFE<br>FYHKCDNTCMESVKNGTYDYPKYSEEAKL<br>NREKIDGSHHHHHH |

**Supplementary Table 3: GISAID strain accession codes**

| Strain | EPI_ISL_ID |
|---|---|
| A/Victoria/4897/2022 (H1N1)pdm09-like virus | EPI_ISL_16714268 |
| A/Thailand/8/2022 (H3N2)-like virus | EPI_ISL_16014504 |
| A/Wisconsin/67/2022 (H1N1)pdm09-like virus | EPI_ISL_15928563 |
| A/Massachusetts/18/2022 (H3N2)-like virus | EPI_ISL_13897304 |
| A_/_H5N1\|A/Cambodia/i0125001G/2024 | EPI_ISL_18823967 |
| A_/_H5N1\|A/California/155/2024 | EPI_ISL_19512043 |
| A_/_H5N1\|A/Washington/239/2024 | EPI_ISL_19512045 |
| A_/_H5N1\|A/California/146/2024 | EPI_ISL_19473581 |
| A_/_H5N1\|A/California/168/2024 | EPI_ISL_19512044 |
| A_/_H5N1\|A/Washington/240/2024 | EPI_ISL_19512046 |
| A_/_H5N1\|A/Washington/239/2024 | EPI_ISL_19531298 |
| A_/_H5N1\|A/Texas/37/2024 | EPI_ISL_19027114 |
| A_/_H5N1\|A/Washington/240/2024 | EPI_ISL_19531299 |
| A_/_H5N1\|A/California/173/2024 | EPI_ISL_19531296 |
| A_/_H5N1\|A/Cambodia/SVH240441/2024 | EPI_ISL_19312044 |
| A_/_H5N1\|A/Washington/253/2024 | EPI_ISL_19531302 |
| A_/_H5N1\|A/Cambodia/KSH240409/2024 | EPI_ISL_19353003 |
| A_/_H5N1\|A/Colorado/134/2024 | EPI_ISL_19280426 |
| A_/_H5N1\|A/Washington/254/2024 | EPI_ISL_19531303 |
| A_/_H5N1\|A/Cambodia/24070331/2024 | EPI_ISL_19312043 |
| A_/_H5N1\|A/British_Columbia/PHL-2032/2024 | EPI_ISL_19548836 |
| A_/_H5N1\|A/Washington/251/2024 | EPI_ISL_19531300 |
| A_/_H5N1\|A/Washington/252/2024 | EPI_ISL_19531301 |
| A_/_H5N1\|A/Colorado/139/2024 | EPI_ISL_19294964 |
| A_/_H5N1\|A/Washington/255/2024 | EPI_ISL_19552697 |
| A_/_H5N1\|A/California/152/2024 | EPI_ISL_19497981 |
| A_/_H5N1\|A/Colorado/138/2024 | EPI_ISL_19294962 |
| A_/_H5N1\|A/California/153/2024 | EPI_ISL_19497980 |
| A_/_H5N1\|A/Colorado/137/2024 | EPI_ISL_19294963 |
| A_/_H5N1\|A/Colorado/109/2024 | EPI_ISL_19263923 |
| A_/_H5N1\|A/Michigan/90/2024 | EPI_ISL_19162802 |
| A_/_H5N1\|A/California/150/2024 | EPI_ISL_19497982 |
| A_/_H5N1\|A/Victoria/149/2024 | EPI_ISL_19156871 |
| A_/_H5N1\|A/Cambodia/NIPH-2402155/2024 | EPI_ISL_18879683 |
| A_/_H5N1\|A/Vietnam/KhanhhoaRV1-005/2024 | EPI_ISL_19031556 |
| A_/_H5N1\|A/Cambodia/24020179/2024 | EPI_ISL_19270607 |

| | |
|---|---|
| A_/_H5N1\|A/California/135/2024 | EPI_ISL_19463618 |
| A_/_H5N1\|A/California/134/2024 | EPI_ISL_19463619 |
| A_/_H5N1\|A/Cambodia/24020155/2024 | EPI_ISL_19270605 |
| A_/_H5N1\|A/California/150/2024 | EPI_ISL_19544645 |
| A_/_H5N1\|A/California/147/2024 | EPI_ISL_19481947 |
| A_/_H5N1\|A/California/171/2024 | EPI_ISL_19531294 |
| A_/_H5N1\|A/California/149/2024 | EPI_ISL_19481949 |
| A_/_H5N1\|A/Khanh_Hoa/RV1-005/2024 | EPI_ISL_19000405 |
| A_/_H5N1\|A/California/172/2024 | EPI_ISL_19531295 |
| A_/_H5N1\|A/California/148/2024 | EPI_ISL_19481948 |
| A_/_H5N1\|A/Michigan/91/2024 | EPI_ISL_19177746 |
| A_/_H5N1\|A/California/151/2024 | EPI_ISL_19531293 |
| A_/_H5N1\|A/Missouri/121/2024 | EPI_ISL_19413343 |
| A_/_H5N1\|A/Washington/UW32495/2024 | EPI_ISL_19541397 |
